# Awakening intracellular immunity for functional HIV cure

**DOI:** 10.64898/2026.01.04.697570

**Authors:** Alessio Lanna, Federica Rinaldi, Christof Stingone

## Abstract

HIV persists in long-lived CD4⁺ T cell reservoirs despite antiretroviral therapy (ART)^1^. Elite controllers suppress viraemia without ART^2^, yet reservoir reactivation emerges with immune ageing^3^. Here we show that transient reprogramming of patient-derived CD4⁺ T cells restores their ability to eliminate HIV-infected reservoirs, excising integrated proviral DNA. A defined compound, or physiological induction, drove rapid reprogramming ex vivo, enabling clearance of HIV DNA within hours to days of treatment, independently of ART. Single-cell RNA sequencing revealed activation of an antiviral telomere transfer programme^4^ that exceeds elite-like control. In humanised mice, adoptive transfer of reprogrammed patient CD4⁺ T cells, or *in vivo* reprogramming of murine T cells, eliminated HIV DNA across reservoirs, with undetectable viral genomes persisting for months. Modelling predicted that residual proviral reactivation would be governed by rare stochastic events, rendering viral rebound unlikely within a human lifespan. These findings identify a previously unrecognised form of intracellular immunity and establish a defined route to a functional HIV cure arising from the CD4⁺ T cell itself.

## INTRODUCTION

HIV remains incurable despite suppressive antiretroviral therapy (ART) because latent reservoirs within long-lived CD4⁺ memory T cells persist for life^1,5^. Even rare elite controllers, who naturally restrain viral replication, eventually experience viral rebound as immune ageing erodes control^2,6^. These observations suggest the existence of an unrecognised link between HIV control and age-related T cell dysfunction.

HIV infection itself accelerates immunosenescence and senescent-like features in non-senescent cells^7,8,9^, yet whether rejuvenating CD4⁺ T cells alone is sufficient to eliminate integrated HIV reservoirs is unknown.

We recently identified an intercellular telomere-transfer pathway that rejuvenates T cells^4,10^, induces stem-like states and reverses senescence by dismantling sestrin–MAP kinase stress complexes (sMACs)^4,10–12^. Correspondingly, sMAC disruption restores telomere transfer^13^, reprogramming T cell responses through feed-forward rejuvenation cascades^4,10–13^. We asked whether such rejuvenation could reactivate intrinsic antiviral functions in CD4⁺ T cells to eradicate HIV reservoirs. We show that T cell rejuvenation awakens a previously unrecognised intracellular mechanism that clears HIV DNA without ART, providing a defined cellular route to a functional HIV cure.

To probe the impact of T cell rejuvenation on HIV latency, we infected primary human CD4⁺ T cells with lentiviral vectors^11^ encoding GFP to mark infected cells. Pharmacological disruptors of sestrin–MAPK complexes (DOS), which induce T cell rejuvenation^13^, resulted in near-complete loss of integrated HIV DNA after seven days of culture (qPCR; **Fig. 1a**). Clearance correlated with dismantling of sestrin–MAPK signalling (**Fig. 1b**).

**Figure 1:**
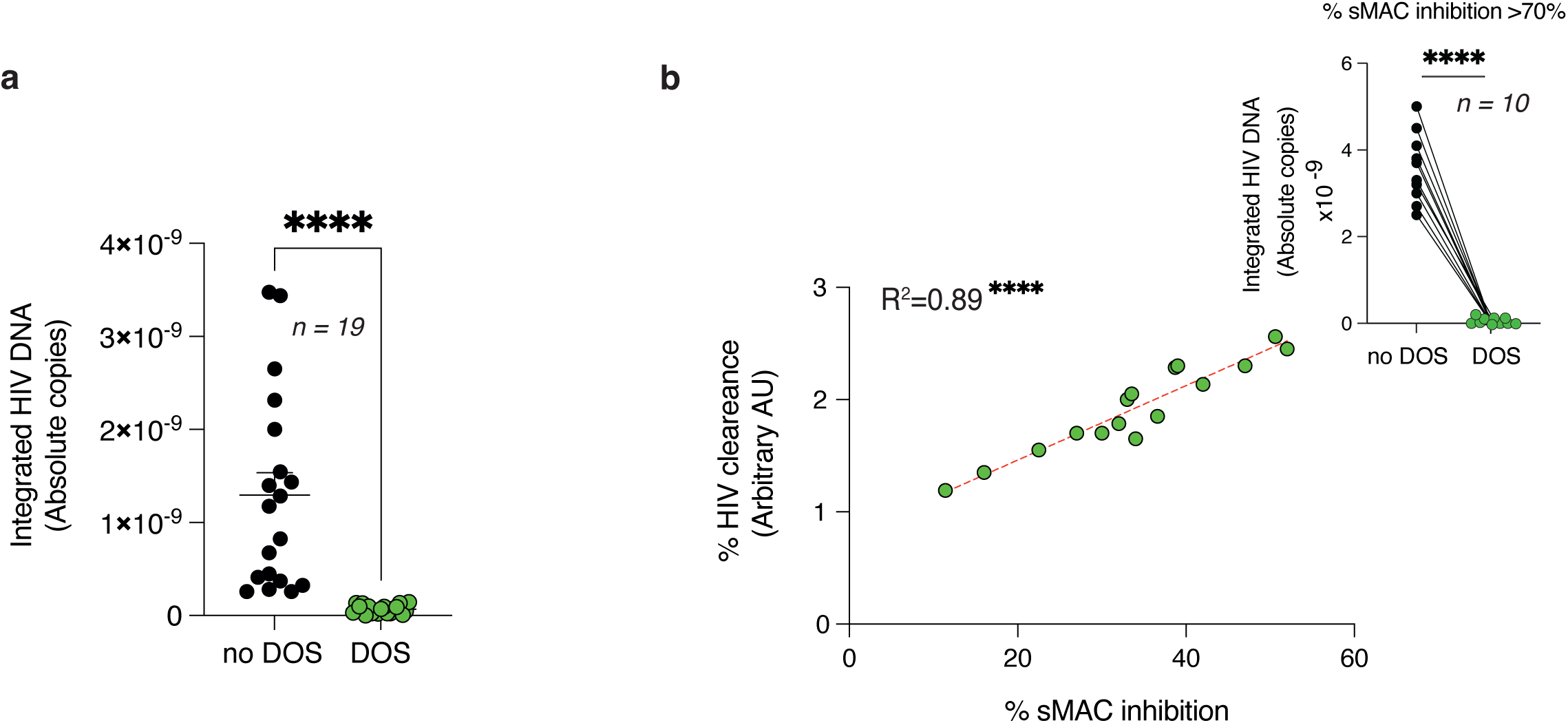
Rejuvenated CD4^+^ T cells cure HIV. **a**, Real time PCR in human CD4^+^ T cells (*n* = 19 donors) infected with HIV lentiviral vectors for 72 hours then treated or not with DOS (4 cycles, dose escalation over one week). Note that DOS treatment eradicates proviral HIV from T cell DNA. **b**, Correlation between HIV clearance as determined via qPCR and Sestrin-MAPKinase (sMAC) disruption (4h single dose treatment) measured by intracellular flow-cytometry among infected human CD4^+^ T cells. Data are from *n* = 16 donors. Note that in some donors (∼40%) with elevated sMAC inhibition, HIV is cleared after a single 4-hour exposure to DOS (*n* = 10 donors). In **a**, **b** (**top**) a paired Student’s t-test and in **b**, Pearson’s correlation coefficient. Data are shown as mean ± s.e.m. **** P < 0.0001. R² = 0.89.

Notably, in approximately 40% of donors, elevated sMAC levels rendered T cells more susceptible to DOS-induced sMAC disruption^11–13^, enabling rapid eradication of HIV proviral DNA within 4 hours of treatment (**Fig. 1b, top right**). Thus, sMAC disruption appears sufficient to eliminate HIV proviral DNA from infected T cells.

To define underlying mechanisms, we performed single-cell RNA sequencing on GFP⁺ HIV-infected CD4⁺ T cells. This enabled isolation of *bona fide* infected cells and comparison of transcriptional pathways between T cells that retained or had purged HIV DNA (**Fig. 2a**). Single-cell RNA sequencing of rejuvenated, GFP-tracked HIV-cleared CD4⁺ T cells revealed telomere transfer^4^ transcriptional signatures enriched for homologous recombination, vesicle communication, WNT and DNA-repair pathways, especially among effector-like T cells where the sMAC is most abundant^4,11–13^ (**Fig. 2b**). Several downstream effectors of telomere transfer, including TZAP-like factors (ZNF580)^4^ and SERPINE2^10^ were also detected. Gene-set enrichment analysis further showed concordance between rejuvenated, HIV-purged CD4⁺ T cells and immune cells from elite-like individuals^14–16^(**Fig. 2c**), with shared activation of DNA-repair, immune-activation and vesicle-communication pathways. Thus, CD4⁺ T cells that intracellularly clear HIV after rejuvenation adopt transcriptional signatures characteristic of telomere-transfer–induced resilience and of naturally occurring elite-like controllers.

**Figure 2:**
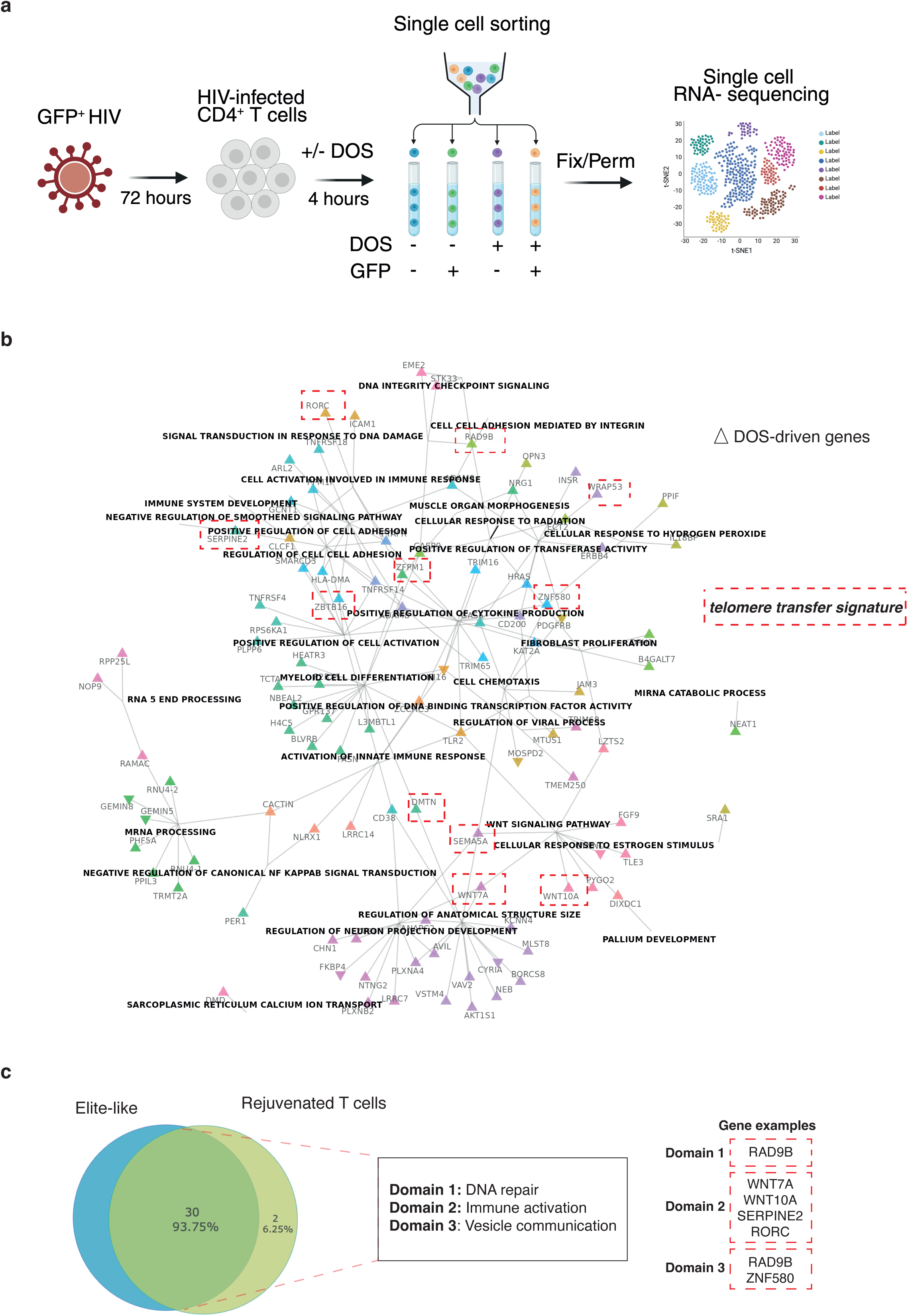
Single cell RNA transcriptomics of HIV functional cure. **a**, Experimental design. Human CD4^+^ T cells were infected with HIV lentiviral vectors encoding a GFP marker gene. The cells were exposed or not to DOS for 4 hours then subjected to single cell sorting to distinguish GFP effectively infected T Cells and GFP negative cells that were not infected, with or without DOS. Following single cell sorting the T cells were then subjected to single cell RNA sequencing. This allowed depicting transcriptomic landscape triggering HIV eradication. **b**, UMAP derived then dot plots depicting gene expression comparison between effectively HIV infected CD4^+^ T cells (GFP^+^) that have been cured or not with DOS, focused on senescent T cell effectors with the highest sMAC levels^11–13^. **c**, David guided gene expression profiles further demonstrated similarities between rejuvenated T cells that have been cured from HIV and naturally occurring elite-like humans. Note that the telomere transfer signature characterised by DNA repair, homologous recombination, vesicle transfer and stem-like related WNT signalling among the HIV cured T cells. HIV DNA removal was confirmed in parallel by genomic PCR. Data are from *n* = 21 donors.

We reasoned that T cell rejuvenation might reactivate an intracellular immunity pathway that competes with HIV integrase for access to host DNA. A candidate mechanism was the RAG recombinase system^17^, which shares structural homology with HIV integrase^18–19^ and is reactivated upon T cell reprogramming through WNT signalling^13,20^, consistent with single-cell transcriptomic signatures and elite-like human profiles.

We therefore tested whether RAG activity is required for HIV clearance. HIV proviral DNA could not be eliminated from RAG-deficient CD4⁺ T cells generated by lentiviral shRAG transduction (qPCR; **Fig. 3a**). To define the mechanism, we focused on BRD9, a chromatin-relaxation factor required for HIV integration^21^ and reported to interact with RAG proteins^22,23^. Chromatin immunoprecipitates from HIV-infected CD4⁺ T cells prepared with anti-integrase antibodies were incubated with recombinant RAG proteins, followed by BRD9 pulldown and integrase detection. Recombinant RAG displaced HIV integrase from BRD9-containing chromatin (**Fig. 3b**) and eliminated proviral DNA within hours, in a BRD9-dependent way (**Fig. 3c**). Correspondingly, rejuvenated HIV-cleared CD4⁺ T cells contained RAG–BRD9 complexes devoid of HIV integrase (**Fig. 3d**). Of note, RAG and HIV integrase expression were strictly transient, becoming undetectable within 72 hours of rejuvenation treatment and cleared through proteasomal degradation (**Fig. 3e** and Extended Data Fig. 1a). Likewise, the reversible appearance of DNA-damage foci^10–13^, and the induction of a stem-like state^13^, confirmed that this pathogen-dismantling machinery is both safe and transient while not inducing cell death (Extended Data Fig. 1b-d). Thus, CD4^+^ T cell rejuvenation activates a RAG-based intracellular immunity pathway that eradicates integrated HIV DNA (**Fig. 3f, proposed model**).

**Figure 3:**
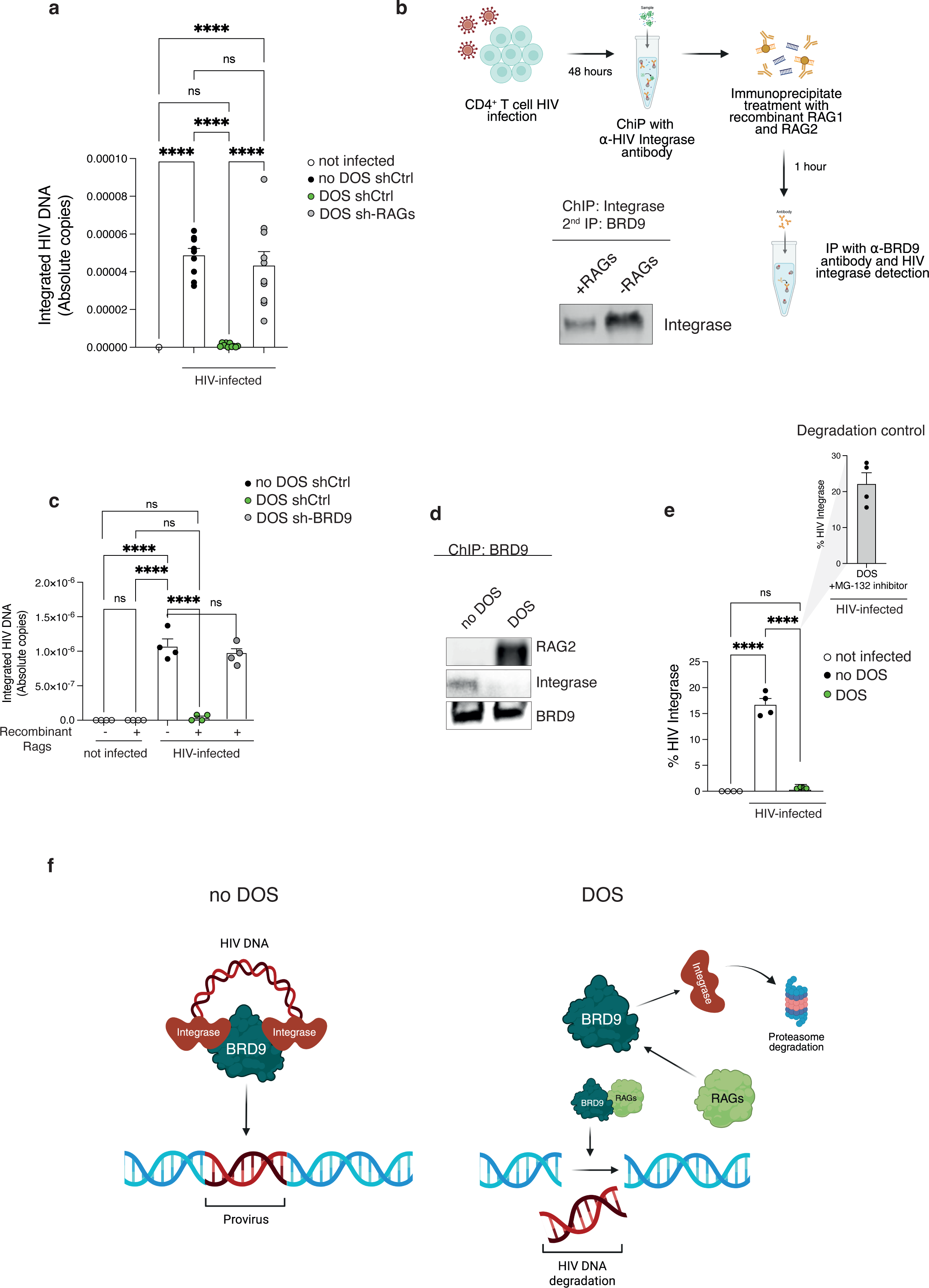
Rejuvenated CD4^+^ T cells cure HIV via RAG proteins. **a**, Real time PCR in human CD4^+^ T cells infected with HIV-1 lentiviral vectors for 72-hours then treated or not with DOS for 4 hours. Note that DOS treatment eradicates HIV only in RAG expressing T cells (shCtrl; *n* = 7 donors) but not in RAG depleted T cells that have been infected with shRAG (*n* = 10 donors), demonstrating RAG dependency. **b**, Effect of recombinant RAG proteins on HIV integrase. Human CD4 T cells infected with HIV lentiviral vectors were subjected to immunoprecipitation (IP) with anti-HIV integrase. The IP was then reacted with recombinant RAG proteins (1:1 ratio) for 1 hour followed by further IP with anti BRD-9 and anti-HIV integrase detection. Note that RAG protein treatment directly breaks BRD-9 HIV integrase binding. Representative of *n* = 4 experiments (4 donors). **c**, Effect of recombinant RAG proteins on HIV DNA. Human CD4^+^ T cells were infected with shCtrl or shBRD9 HIV lentiviral vectors and DNA extracted 72 hours later. The extracted DNA was then incubated for 4 hours with recombinant RAG1 and RAG2 proteins (1:1 ratio), followed by a subsequent DNA purification. HIV DNA was detected by Real time PCR (*n* = 4 donors). **d**, Human CD4^+^ T cells infected with HIV lentiviral vectors were treated or not with DOS for 4 hours. The T cells were then subjected to IP with anti BRD-9 followed by immunoblotting, as indicated. Note that DOS treatment phenocopies recombinant RAGs. Representative of *n* = 4 experiments (4 donors). **e**, Human CD4+ T cells infected with HIV lentiviral vectors were treated or not with DOS for 4 hours and the expression of HIV integrase was measured by intracellular flow-cytometry among the GFP positive cells. In some cultures, the infected T cells were pretreated with the proteasome inhibitor MG-132 (10uM) 30 minutes prior to DOS treatment (n = 4 donors). Note that DOS treatment triggers HIV integrase proteasome degradation. Similar integrase resolution was observed following repeated (dose escalation) DOS treatment in one-week cultures. **f**, Model of the functional HIV cure. T cell rejuvenation triggers RAG protein expression (controlled by the WNT elite-like pathway, to outcompete HIV integrase for BRD-9 binding). This results in both integrase displacement from infected DNA and RAG mediated viral degradation. Subsequently, both HIV integrase and RAG proteins are degraded via proteasome, safeguarding genomic stability. In **a**, **c**, **e** one-way ANOVA with Bonferroni post hoc correction. Data are shown as mean ± s.e.m. **** P < 0.0001; ns, not significant.

This clearance was reproducible in *ex vivo* CD4⁺ T cells from HIV⁺ donors (*n* = 61), regardless of ART status, following pharmacological or telomere vesicle rejuvenation during a one-week culture (**Fig. 4a**). In these same experiments, a subset of patient T cells with elevated sMAC burden underwent near-complete proviral clearance even after a single 4-hour rejuvenation cycle, consistent with an infection cure model triggered by sMAC disruption.

**Figure 4:**
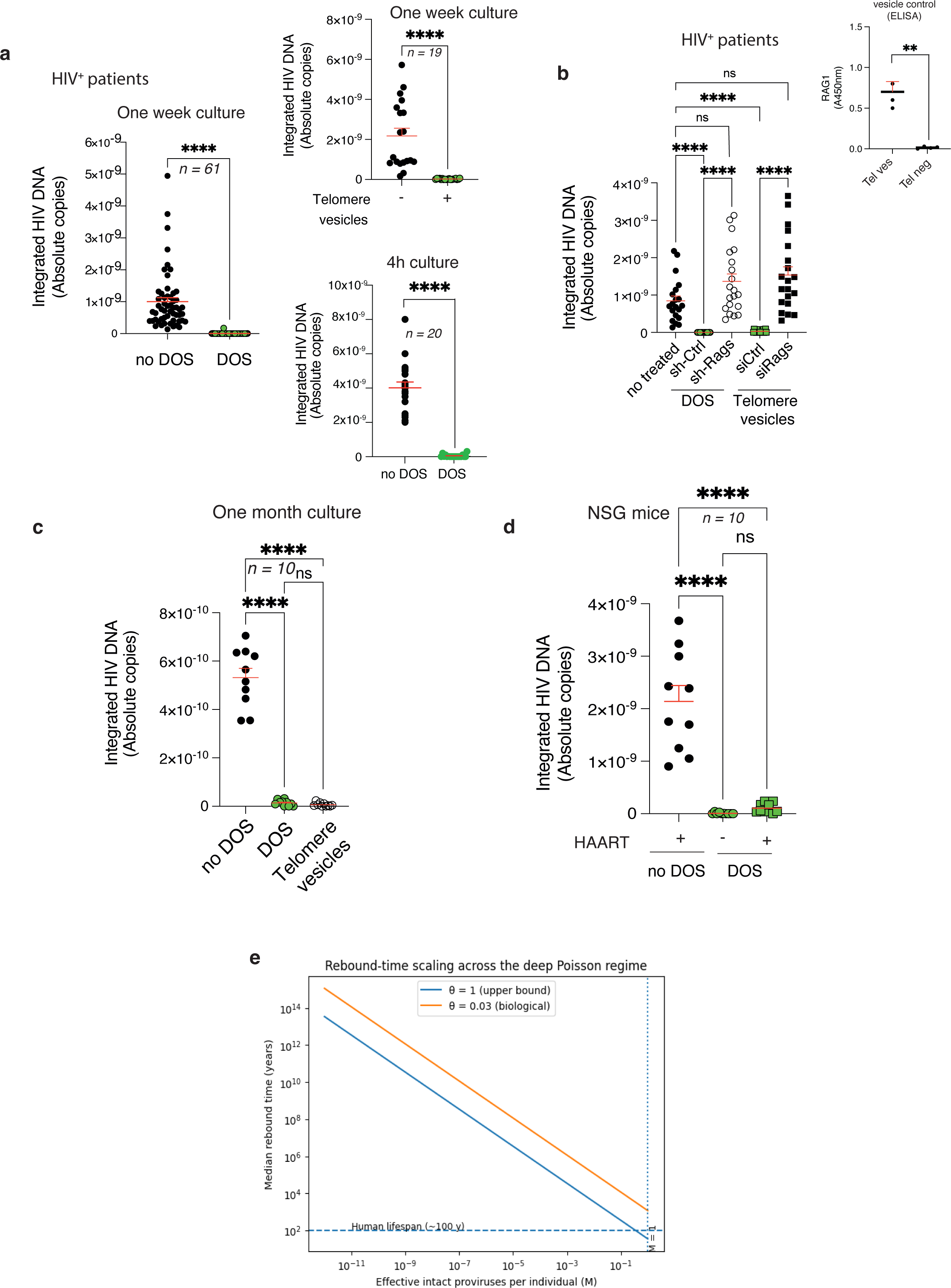
Rejuvenation cures HIV in patient CD4^+^ T cells, *in vivo*. **a**, Sixty-one HIV infected patients were recruited and their CD4^+^ T cells exposed or not to DOS for 1 week (four subsequent, dose escalation treatments). The T cell DNA was then assessed for presence of integrated HIV DNA (proviruses) by real time PCR. Note that similar to lentiviral vector models, DOS-driven T cell rejuvenation eradicates integrated HIV resulting in functional cure. In parallel, telomere vesicle treatment was also assessed (**top right**). Note that in a subset of patients, HIV clearance occurs within 4h of single DOS treatment (*n*= 20 donors; **bottom right**). **b**, Effect of T cell rejuvenation by DOS or telomere vesicles on HIV^+^ patient T cells is RAG dependent. T cells were either depleted or not of RAG proteins prior to DOS; or subjected to transfer of autologous APC telomere vesicles carrying RAG proteins or not, as indicated (*n*= 20 donors). Note that T cell rejuvenation cures HIV independently of ART regimens but requires T cell RAGs or telomere vesicles carrying RAG proteins. RAG content in telomere vesicles (Tel ves) was confirmed by ELISA upon extraction from autologous APCs^4^ (*n*= 4 donors; **top right**). Tel neg, APC telomere negative vesicles. **c**, The long-lasting cure was confirmed in one month cell cultures, without any antiretroviral treatments (ART), after DOS or telomere vesicle addition (*n*= 10 donors). Cells were cultured with DOS or telomere vesicles for one week, followed by treatment interruption and further activation with anti CD3/CD28 plus rhIL2 every ten days. **d**, CD4^+^ T cells from patients were transferred into nude NSG mice treated with or without DOS weekly (four DOS cycles; as detailed in Methods). The T cells were then assessed by real time PCR from the blood of the NGS recipients five weeks later. In some experiments, the animals were subjected to daily ART, as indicated. Real time PCR data of patient CD4^+^ T cells derived from NSG mice is shown (*n* = 10 mice per group). **e**, Rebound time scaling under conservative and illustrative assumptions. The black line shows the conservative upper-bound model assuming all residual proviral DNA is reactivatable (θ = 1). The green dashed line shows an illustrative sensitivity incorporating limited proviral reactivation (θ = 0.03), shown for context. The dotted horizontal line indicates a human lifespan (∼100 years). In **a**, a paired Student’s t-test; in **b**-**d**, one-way ANOVA with Bonferroni post hoc correction. Data are shown as mean ± s.e.m. **** P < 0.0001; ns, not significant.

Importantly, as per DOS treatment, telomere-vesicle transfer induced similar RAG-dependent HIV clearance in patient T cells but only when vesicles carried RAG proteins, indicating a shared, RAG-driven rejuvenation clearance mechanism (**Fig. 4b**). Clearance was validated by HIV mRNA quantification (Extended Data Fig. 2a) and remained durable for over four weeks even after withdrawal of either rejuvenation treatment and antiretrovirals, and after subsequent expansion of the cells upon activation with anti-CD3/CD28 plus IL-2 (**Fig. 4c**). These clearance data were further confirmed by viral outgrowth assays^24^ and by intact proviral DNA measurements using both IPDA^25^ and Alu-PCR^26^ (Extended Data Fig. 2b-d). Moreover, T cells from HIV⁺ donors that had reached the HIV-proviral-undetectable state expressed key signalling molecules previously identified by single-cell transcriptomics, as demonstrated by intracellular flow cytometry (Extended Data Fig. 3). Together, these findings demonstrate that HIV can be eradicated from patients T cells within days of rejuvenation.

To assess HIV eradication *in vivo*, NSG mice received HIV⁺ donor CD4^+^ T cells then subjected to weekly *in vivo* purge treatments with sMAC disruptors (DOS). Strikingly, qPCR showed absence of integrated HIV among circulating CD4^+^T cells 5 weeks later, in contrast to persistent infection in ART treated controls (**Fig. 4d**). We also confirmed persistent HIV clearance in immune challenged CD45.1/2 donor-recipient systems, including brain and gut reservoirs where ART penetration is limited, up to one year after DOS interruption (Extended Data Fig. 4). Consistently, mathematical modelling predicted that reactivation of a single residual provirus, if present, would exceed a human lifespan (**Fig. 4e**). T cell rejuvenation enables sustained HIV clearance *in vivo*.

We identify a previously unrecognised form of intracellular pathogen dismantlement, enabling a functional HIV cure. Functional cure required transient licensing of ancestral RAG recombinases, which are normally silenced in mature T cell lineages^27^ but become briefly available upon T cell rejuvenation. Importantly, RAG and HIV integrase expression were strictly transient and preserved genomic stability.

Functional cure did not require ART and was integration sequence agnostic, since it relied on RAG binding to specific CD4^+^ T cell chromatin factors, such as BRD9. Thus, through host chromatin interactors, functional cure operates independently of integration site identity, with no requirement for proviral sequence mapping or specific mutational landscapes. While functional cure occurred independently of ART, it remains to be determined whether clinical HIV integrase inhibitors that often display off target RAG inhibition^18–19,28,29^ be compatible or interfering with RAG driven curative approaches.

In contrast to elite controllers - who suppress viraemia yet remain unable to eliminate latent reservoirs^25^ - reprogrammed T cells directly targeted HIV integrase bound chromatin, initiating targeted excision of proviral DNA. Moreover, elite controllers preserve WNT-driven stem-like T cell pools but cannot overcome stress-mediated repression of RAGs^30,31^; and high thymic output in elite controllers further reinforces peripheral RAG silencing and recombinase inhibition^32^

Reprogramming overcame these barriers and enabled RAG-mediated displacement of HIV integrase and excision of proviral DNA. These strategies circumvent CRISPR-based limitations^33^, which require prior knowledge of HIV integration sites, and avoid reliance on “shock-and-kill” regimens that are undermined by viral escape and incomplete cytolytic clearance^34^, or “lock-in” strategies that do not clear integrated reservoir^35^. Together, our results indicate that HIV mutation burden is not a barrier to achieving a functional cure.

Instead, competitive displacement of the HIV integrase interactome on host chromatin is sufficient to trigger rapid proviral excision and reservoir elimination. The sequence-independent nature of this pathway suggests a general curative principle against other chronic conditions sustained by integrated pathogens or retroelements.

Across biological and mathematical settings therefore, functional cure consistently reduced HIV reservoirs to undetectable state, preventing viral resurgence under rebound-permissive conditions, and constraining stochastic rebound trajectories beyond human lifespan. While it is not possible to formally establish absolute sterilisation, these results may be compatible with durable treatment free HIV negative state, similar to rare outcomes observed following bone marrow transplant in people living with HIV.

## Data availability

The data generated or analysed in this study are included in the manuscript, and its Extended data Figures.

## Acknowledgments.

We are grateful to all the blood donors who made this work possible and Alessandra Latini for C. S. support. This work was funded by Sentcell ltd. The funder had no role in study design or decision to publish the manuscript. A.L. is Full Professor of the University College London and the Chief Executive Officer of Sentcell ltd.

## Contributions

A.L. discovered the functional HIV cure, directed the study, analysed the data and wrote the paper; F.R. performed and analysed experiments; C. S. provided HIV^+^ clinical samples and discussed with A.L.

## Competing interests

A.L. is a shareholder to Sentcell ltd and the sole inventor of the DOS pharmaceutics and further methods to cure HIV where Sentcell ltd figures as the Applicant. A.L. and F.R. are supported by Sentcell ltd.

---

**Extended Data Fig. 1:**
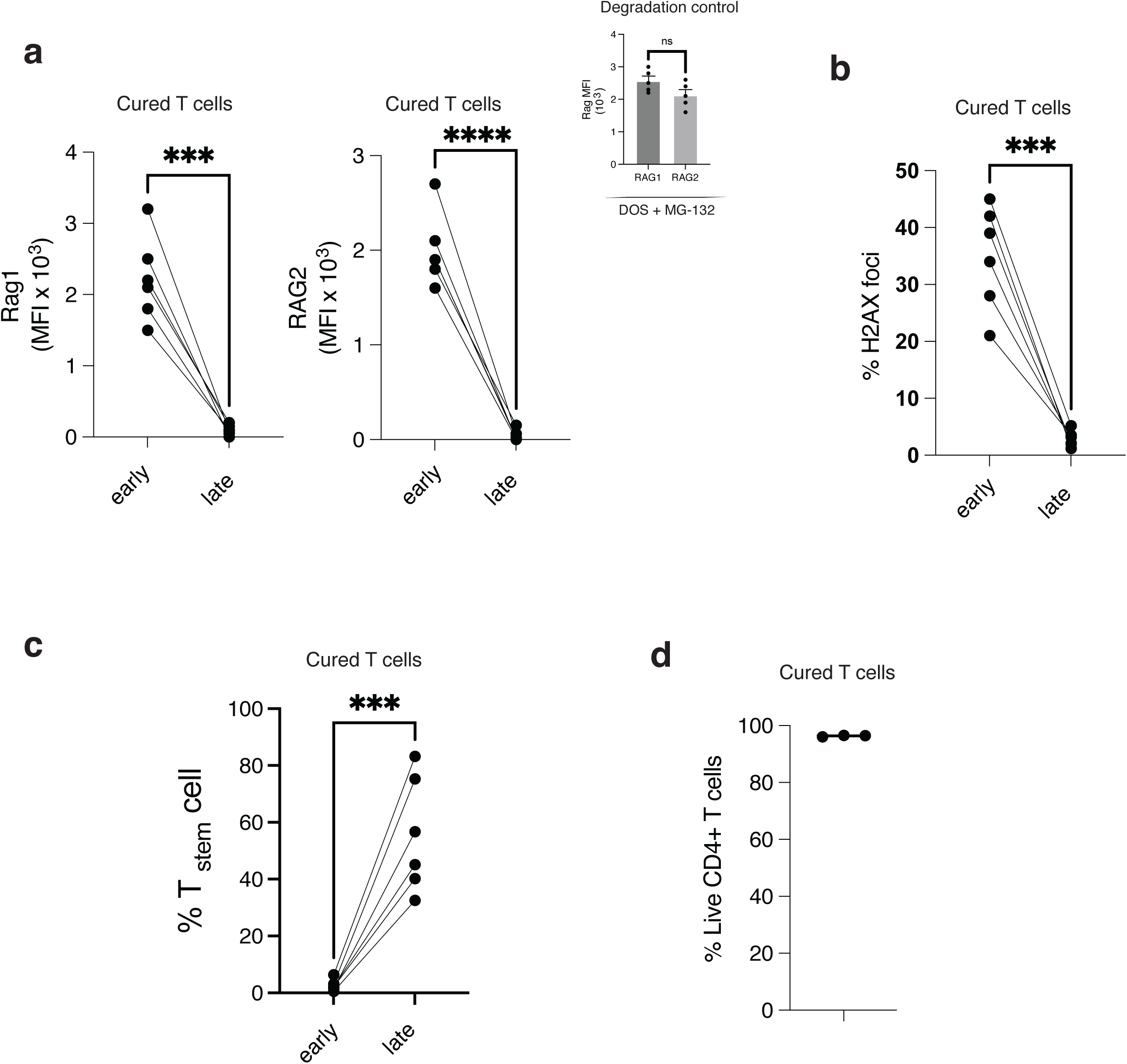
Transient RAG licensing, reversible DNA-damage, and preserved cell viability in HIV cured T cells. **a**, RAG1/RAG2 expression in HIV-infected human CD4⁺ T cells following DOS treatment (single dose, 4h), showing rapid induction and complete resolution within 72 hours of treatment. MFI, mean fluorescence intensity (RAG1, *n* = 6 donors; RAG2, *n* = 5 donors). In some experiments, the T cells were pre-treated with MG-132 (1μM) prior to DOS treatment (**top right**). **b**, Reversible DNA-damage signalling assessed by γH2AX foci formation following rejuvenation driven HIV eradication, with resolution over time (96h) and no evidence of persistent genomic stress (*n* = 6 donors). **c**, Stem-like T cell induction among cured T cells (CD4^+^ CD45RA^+^ CD27^+^ CD28^+^ CCR7^+^, CD62L^+^, CD95^+^) after a one-week culture and repeated dose treatment throughout (dose escalation). Data are from *n* = 6 donors. **d**, Cell viability (Annexin V) show no increase in cell death following DOS exposure, indicating that HIV clearance is not attributable to cytotoxicity or clonal selection (*n* = 3 donors). Similar RAG/ γH2AX resolution was observed following repeated (dose escalation) DOS treatment in one-week cultures. In **a**-**d**, a paired Student’s t-test. Data are shown as mean ± s.e.m. *** P < 0.001; **** P < 0.0001; ns, not significant.

**Extended Data Fig. 2:**
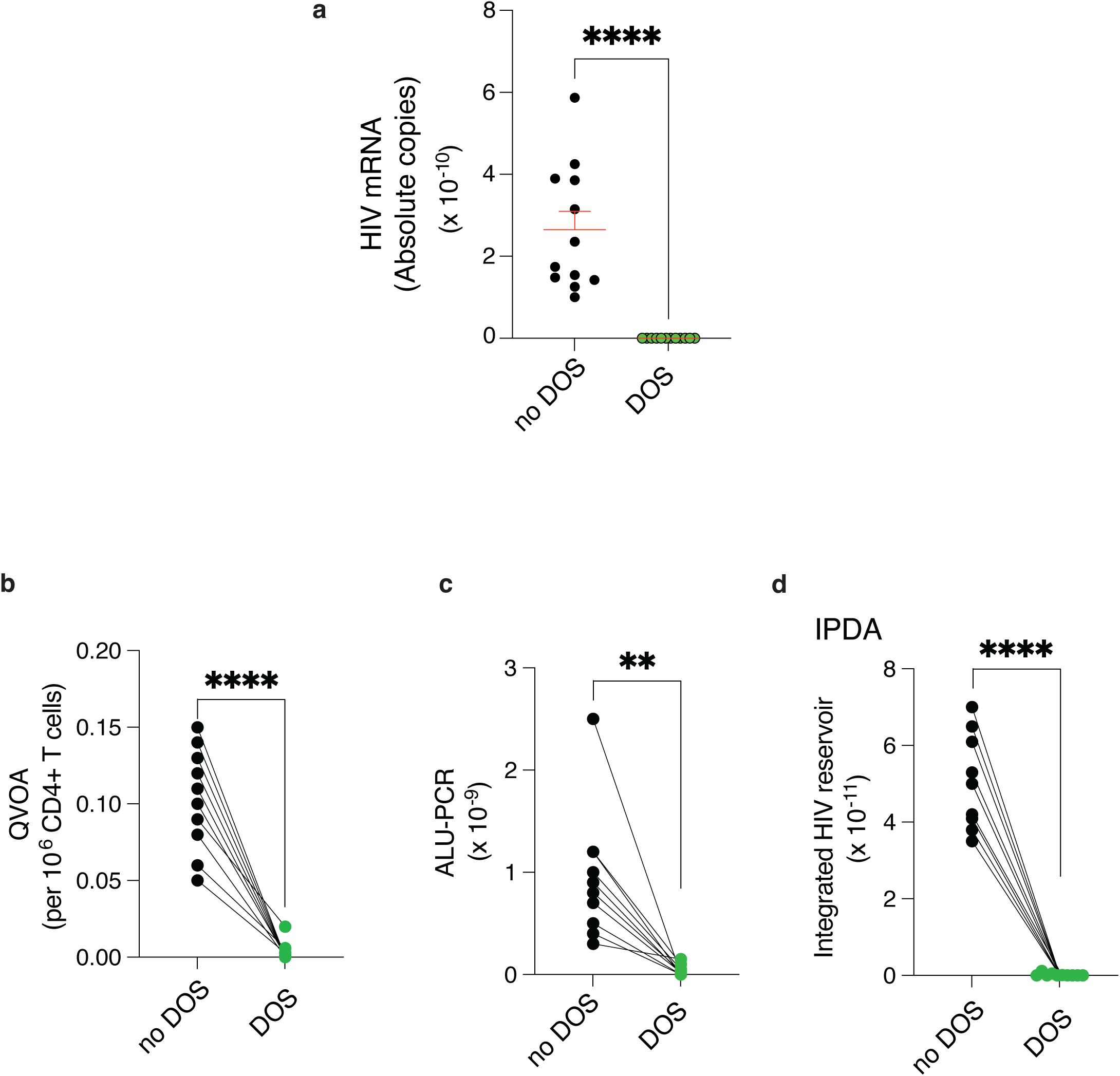
Confirmation of HIV eradication upon T cell rejuvenation. CD4**^+^**T cells from HIV^+^ donors were subjected to DOS rejuvenation treatment (4 cycles, dose escalation over one week) after which HIV mRNA levels were immediately detected (**a;** *n* = 11 donors). HIV reservoir depletion was further assessed by Quantitative viral outgrowth assay (**b**, QVOA), quantification of integrated HIV DNA by Alu–PCR (**c**) and intact proviral DNA assay (**d**; IPDA) demonstrating depletion of genetically intact HIV proviruses. In **a**-**d**, a paired Student’s t-test. Data are shown as mean ± s.e.m. ** P < 0.01; **** P < 0.0001.

**Extended Data Fig. 3:**
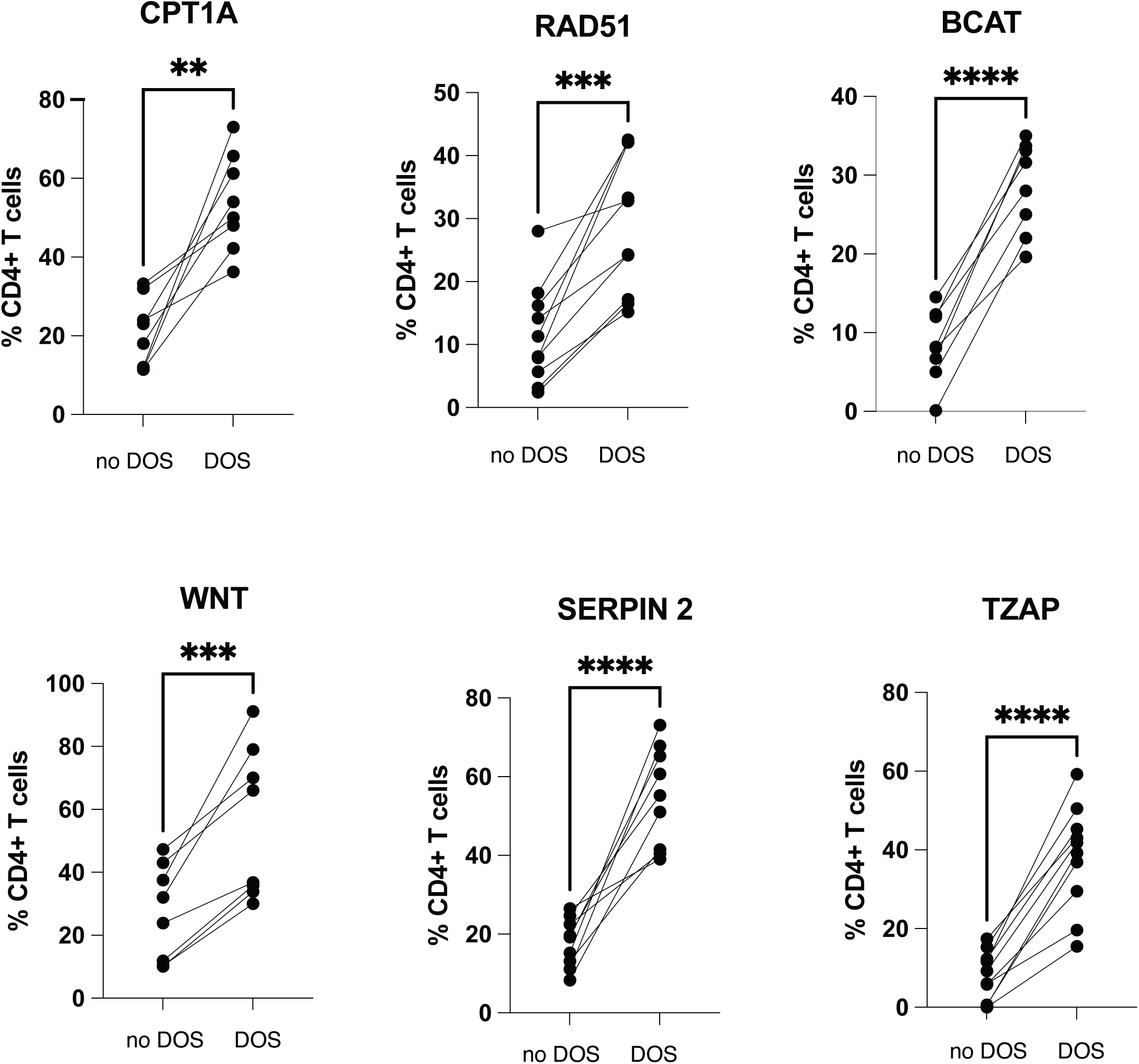
Flow-cytometric validation of single cell transcriptomics. Flow-cytometric analysis of CD4⁺ T cells from HIV⁺ donors that reached an HIV-proviral-undetectable state following DOS treatment and subjected to single cell sequencing. Expression of key signalling molecules/nodes identified by single-cell RNA sequencing is shown, validating transcriptional programmes associated with HIV clearance. Data are representative of *n* = 10 donors. A paired Student’s t-test was used. Data are shown as mean ± s.e.m. ** P < 0.01; *** P < 0.001 **** P < 0.0001.

**Extended Data Fig. 4.**
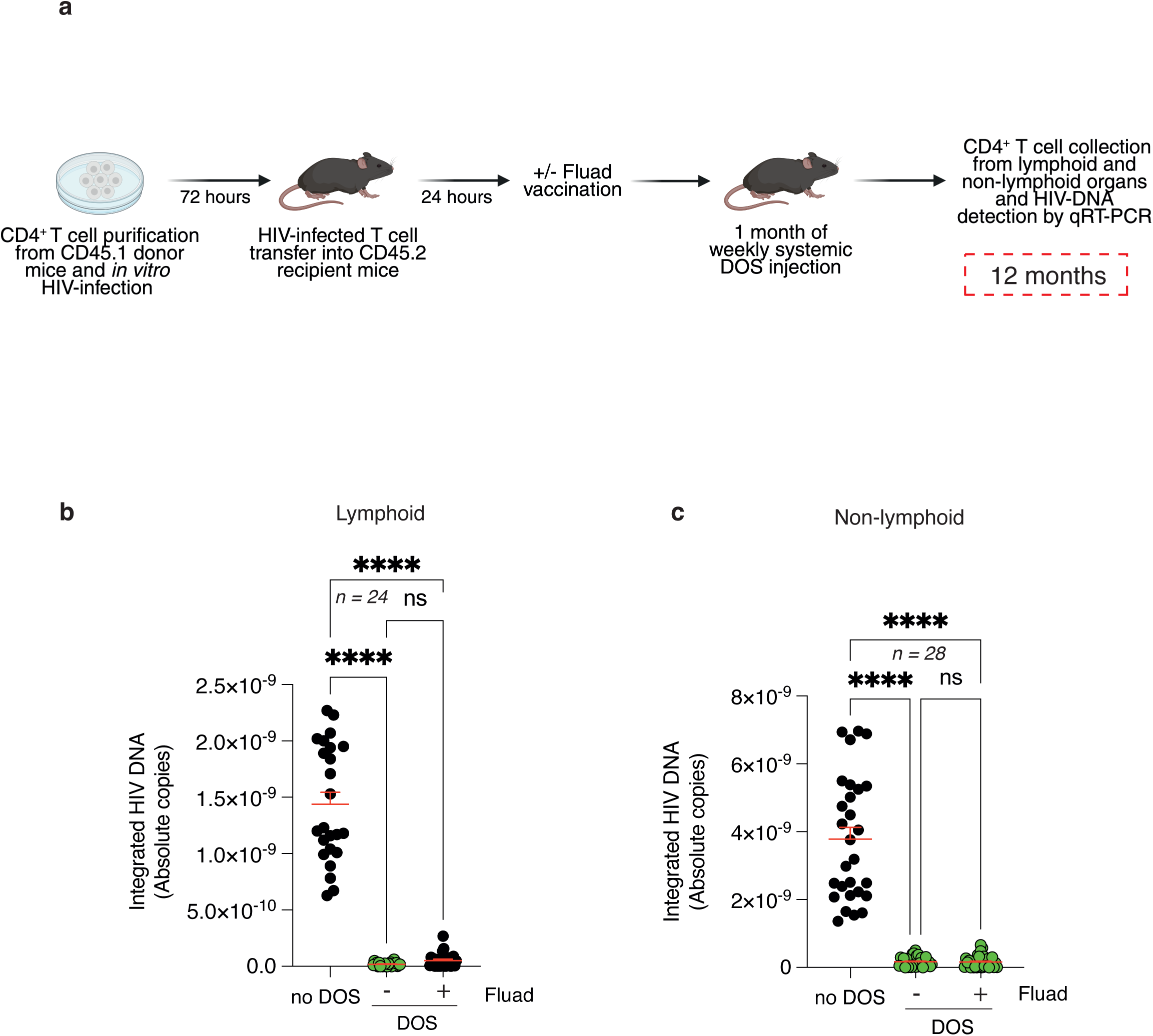
Durable HIV eradication even in virological sanctuaries. **a**, CD4^+^ T cells were derived from CD45.1 donor mice *in vitro*, HIV infected with lentiviral vectors then transferred into CD45.2 congenic recipients. The day after, the animals were vaccinated or not with FLUAD to provide an immune challenge followed by systemic weekly DOS treatment. Twelve months later, the infected CD45.1 CD4 T cells were recovered from the CD45.2 recipient lymphoid (**b**, lymph nodes and spleens) and non-lymphoid (**c**, brain and gut) organs and assessed by real time PCR for HIV integration. Note that DOS treatment eradicates HIV even in non-lymphoid organs that are virological sanctuaries. Data are from *n* = 24 animals (lymphoid organs) or *n* = 27 animals (non-lymphoid organs). In, **b**-**c,** one-way ANOVA with Bonferroni post hoc correction. Data are shown as mean ± s.e.m. **** P < 0.0001; ns, not significant.

## Materials and Methods

### Lentiviral vector production

Lentiviral particles were generated in HEK293 cells using standard transient co-transfection as previously described^1^.

### HIV proviral DNA integration assays

CD4⁺ T cells were purified from PBMCs of healthy volunteers upon informed consent and ethical approval (Prot n 26604), using CD4⁺ T cell isolation kit (Miltenyi Biotec, 130-096-533). Cells (2 x 10^5^ in technical triplicates) were preincubated with polybrene (8 μg/mL, 30 min) and infected with scrambled shRNA lentivirus, shRAG1/2 or shBRD9, as indicated (MOI 10; 1×10⁵ cells; Origene). After 72 h, cells were treated with the DOS compound DOS46L starting at 400 nM for 4 hour-pulse (single or sequential dosing, as indicated in legends) or left untreated. Repeated doses were as follows: 400 nm; 1.2 μM; 3μM; 10μM. For telomere vesicle treatment, autologous exosome-like particles were purified as previously described from RAG proficient (shCtrl) or RAG deficient (shRAGs) APCs^2^, then administered at 250 telomere vesicles per T cell culture (4 repeated treatments, every other day). Cells were activated with anti-CD3/CD28 (0.5ug/ml each) and rhIL2 (5ng/mL). Uninfected controls were processed in parallel to establish detection limit (10^-12^) and calling threshold (Ct ≥ 40) throughout. Knock down efficiency was confirmed by qPCR (≥ 80%).

Genomic DNA was extracted using the PureLink Genomic DNA Mini Kit (Invitrogen). Integrated HIV DNA was quantified by qPCR using PowerUp SYBR Green Master Mix (Applied Biosystems) on a QuantStudio 1 system.

HIV-specific (LTR) primers were used:

FW AAATCTCTAGCAGTGGCGCCCGAACAG

RV CCATCTCTCTCCTTCTAGCCTCCGC

and quantification employed the ΔΔCt method. Alternatively, CD4⁺ T cells were isolated from peripheral blood of HIV^+^ volunteers upon informed consent and specific ethical approval (278/ISG/25). For mRNA detection, mRNA suppression post-treatment was confirmed using the same LTR primers following reverse transcription with oligo(dT).

### Additional validation of HIV reservoir depletion

To independently validate HIV reservoir depletion, complementary molecular and functional assays were performed, including quantitative viral outgrowth assays (QVOA), quantification of integrated HIV DNA by Alu–PCR, and the intact proviral DNA assay (IPDA).

#### Quantitative viral outgrowth assay (QVOA)

Replication-competent HIV was assessed using quantitative viral outgrowth assay adapted from established protocols^3,4^. CD4⁺ T cells were isolated from PBMCs by negative selection as above described and subjected or not to one week DOS driven HIV purging. Cell were then plated at limiting dilution in the presence of irradiated allogeneic PBMC feeder cells. Cells were re-activated with PHA (1μg/mL) and cultured in RPMI supplemented with 10% FBS and interleukin-2 (5 ng/mL) for 21 days. Viral outgrowth was monitored by measuring HIV-1 p24 antigen in culture supernatants by ELISA. Wells were scored positive if p24 exceeded background detection thresholds, and infectious units per million (IUPM) CD4⁺ T cells were estimated assuming Poisson-distributed infected cells.

#### Quantification of integrated HIV DNA (Alu–PCR)

Integrated HIV DNA was quantified using a nested Alu–gag PCR assay as previously described^5,6^. Genomic DNA was extracted from purified CD4⁺ T cells. First-round PCR amplified host–virus junctions using an Alu forward primer (5′-GCCTCCCAAAGTGCTGGGATTACAG-3′) and an HIV-1 gag reverse primer (5′-GTTCCTGCTATGTCACTTCC-3′). Nested quantitative PCR was performed using gag-specific primers (forward: 5′-CATGTTTTCAGCATTATCAGAAGGA-3′; reverse: 5′-TGCTTGATGTCCCCCCACT-3′) and probe (5′-FAM-CCACCCCACAAGATTTAAACACCATGCTAA-TAMRA-3′). Results were normalized to a single-copy host gene (ACTB) and reported as integrated HIV DNA copies per 10⁶ CD4⁺ T cells.

#### Intact proviral DNA assay (IPDA)

Genetically intact HIV proviruses were quantified using the intact proviral DNA assay (IPDA) by multiplex droplet digital PCR, as described previously^7^. Genomic DNA was analyzed using primer–probe sets targeting the HIV packaging signal (Ψ) and env, enabling discrimination between intact and defective proviruses. Ψ primers were 5′-TCTCTCGACGCAGGACTCG-3′ (forward) and 5′-TACTGACGCTCTCGCACC-3′ (reverse), with probe 5′-FAM-CTCTCTCCTTCTAGCCTCCGCTAGTCA-BHQ1-3′. env primers were 5′-AGTGGTGCAGAGAGAAAAAAGAGC-3′ (forward) and 5′-GTCTGGCCTGTACCGTCAGC-3′ (reverse), with probe 5′-HEX-CCTTGGGTTCTTGGGAAGTGGG-BHQ1-3′. A single-copy host reference gene (RPP30) was amplified in parallel for normalization. Intact proviruses were defined as droplets positive for both Ψ and env targets. Limiting-dilution and molecular frequency estimates were derived assuming Poisson-distributed infected cells. Samples below the limit of detection were assigned the lower bound of quantification for analysis.

### Flow cytometry

HIV infected CD4⁺ T cells were washed with ice-cold PBS, stained with anti-CD4-APC (Miltenyi, 130-113-222), fixed with BD Cytofix (30 min, 4 °C), and permeabilized with BD Phosflow Perm Buffer III (30 min, on ice). Intracellular HIV integrase was detected using anti-integrase monoclonal antibody (Invitrogen, MA5-18031, 1:100), followed by Alexa Fluor 405 donkey anti-mouse secondary antibody (Invitrogen, A48257, 1:1000). In some experiments, cells were evaluated for H2AX (Cell Signaling, 2595S; 1:100) and stem like reprogramming with antibodies to CD4 (PerCP5.5, BioLegend, 557956; 1:100), CCR7 (Brillant Violet 421, Biolegend, 100540; 1:100); CD27 (PerCP-Vio 700, Miltenyi; 130-120-037; 1:100); CD28 (PerCP/Cy5.5, BioLegend, 302922; 1:100); CD45RA (BV510, BioLegend, 304142; 1:100); CD62L (PE-Vio 770; Miltenyi; 130-113-621; 1:100) and CD95 (FITC, Miltenyi, 130-122-950; 1:100). To confirm single cell pathway transcriptomics, CD4^+^ T cells were further stained with antibodies to CPT1A (Cell Signaling, 657 12252, 1:1000), Rad51 (Abcam, ab63801; 1:100;), Beta-catenin (Cell Signaling, 9562; 1:100), Wnt5 (Cell Signaling, 2392; 1:100), Serpine 2 (Thermofisher, 5016; 1:100) and TZAP (Novusbio, H00003104-B01P,; 1:100). Analysis was performed by flow cytometry.

### Single-cell RNA-sequencing

CD4⁺ T cells were HIV infected and treated with DOS46L as above. GFP⁺ cells were then sorted using a BD FACS Melody. Cells were fixed and permeabilized using Evercode Cell Fixation (Parse Biosciences) and stored at –80 °C.

Library preparation followed the Evercode Whole Transcriptome kit protocol (Parse Biosciences). Reverse-transcribed cDNA incorporated Round-1 for barcoding during reverse transcription with oligo(dT) primers and random hexamers. Each well incorporated a unique well-specific barcode. After pooling, cells were distributed into a Round 2 plate for ligation of a second barcode to the cDNA. A third round of barcoding was then performed in the Round 3 plate, where a final barcode containing the Illumina TruSeq Read 2 sequence and a biotin moiety was added. The resulting samples were split into eight sub-libraries, lysed, and biotinylated cDNA was captured using streptavidin-coated magnetic beads. A template-switching reaction was used to add a 3’ adapter, and the cDNA was then amplified using primers specific to the template-switch adapter and the TruSeq Read 2 sequence (Illumina, San Diego, CA, USA). Amplified cDNA was fragmented, end-repaired, A-tailed, and ligated to the Illumina TruSeq Read 1 adapter. Sub-libraries were assessed for quality with a Fragment Analyzer™ (Agilent Technologies) and quantified by qPCR. Eight sub-libraries were generated, purified on streptavidin magnetic beads, amplified, fragmented, and ligated to Illumina adapters. Library quality was assessed using an Agilent Fragment Analyzer and quantified by qPCR. Samples were sequenced on the Illumina NovaSeq 6000 (100 bp paired-end). Each reaction was performed in 4 technical replicates from *n =* 21 donors.

### scRNA-seq processing and analysis

Sequencing reads were demultiplexed and quality-checked using the Parse Biosciences pipeline (split-pipe v1.5.0). Reads were aligned to the human reference genome (hg38), and cell-by-gene count matrices were generated. Downstream analysis was performed using Seurat (v5.2.1; Hao et al., 2024) in R (v4.4.1). Low-quality cells were excluded based on the following criteria: >10% mitochondrial gene content, <100 or >10,000 detected genes. Gene expression was normalized using centered log-ratio (CLR) transformation.

Cell types were annotated using canonical markers^8^:

- Naïve: CCR7, SELL, LRRN3
- Stem-like: LRRN3, CCR7, SELL, FAS, CXCR3
- Central memory: PASK, ITGB1, VIM, AQP3
- Effector memory RA (EMRA): CCL4, GZMH, GZMA, GNLY, NKG7, CST7
- Effector memory: KLRB1, TNFSF13B, GZMK, CCL5
- Treg: ANXA2, LGALS3, LGALS1, FOXP3, CTLA4, IL2RA

To identify differentially expressed genes (DEGs) between experimental conditions within each manually annotated T cell subtype, the FindMarkers function from the Seurat package was applied. The Wilcoxon rank-sum test was used for statistical testing, and p-values were adjusted for multiple testing using the Benjamini-Hochberg method. Genes with an adjusted p-value < 0.05 were considered significantly differentially expressed. Pathway enrichment analysis was conducted using the Single Cell Pathway Analysis (SCPA) algorithm, in conjunction with the Gene Ontology C5 subcollection (Biological Process, Molecular Function, and Cellular Component categories; https://www.gsea.msigdb.org/gsea/msigdb).

Pathway comparisons were performed between experimental conditions within each T cell subtype. Pathways with a Q-value > 1.14 were considered significantly enriched where Q-value is defined as √[–log10 (Bonferroni-adjusted) p value]. Gene enrichment analysis with public elite like repertoires^15^ was performed by DAVID algorithm.

### Double immunoprecipitation

CD4⁺ T cells (4×10⁷ per reaction) were infected with HIV lentiviral vectors for 72 h. Chromatin was crosslinked and sheared using the SimpleChIP Plus kit (Cell Signaling, #9005). Chromatin was incubated overnight with anti-HIV integrase antibody (Invitrogen, MA5-18031; 1.5 μg/IP), followed by capture with magnetic beads. For RAG competition assays, integrase IP complexes were incubated with recombinant human RAG1 and RAG2 (200 ng each, 1 h, RT). In parallel, BRD9 antibody (Cell Signaling, 58906S; 1.5 μg) was pre-bound to magnetic beads and added to RAG-treated or untreated samples for a second IP (overnight, 4 °C). Complexes were washed, eluted in DTT-containing Laemmli buffer, heated (95 °C, 5 min), and resolved by SDS-PAGE.

### Immunoprecipitation

After 72 h HIV infection, CD4⁺ T cells were treated with DOS46L (400 nm) for 4 h. Chromatin was prepared using the SimpleChIP Enzymatic IP Kit (Cell Signaling, #9003). BRD9 IPs were performed with 1.5 μg BRD9 antibody (58906S) overnight at 4 °C, followed by magnetic bead capture (2 h). Eluted proteins were reverse-crosslinked, resolved by SDS-PAGE, and immunoblotted for HIV integrase (Abcam, ab66645; 1:100), BRD9 (Cell signaling, 71232; 1:100), RAG1 (Cell signaling, 3968S; 1:100) and RAG2 (Invitrogen, PA5-88580; 1:100).

### Mouse studies

Animal approval was received from the Italian Ministry of Health and the UK home office (PP8261278). CD4⁺ T cells from HIV-infected patients were isolated from PBMCs and transferred directly into NOD/SCID/IL2Rγ⁻/⁻ (NSG) mice. In these experiments, patient CD4⁺ T cells were transferred (2 x 10^6^; intra-venous injection) into NSG mice, along with 10^6^ human (autologous) APC feeder cells. After transfer, mice received daily ART (0.1 mg emtricitabine, 4mg rilpivirine, and 2.14mg tenofovir alafenamide) in drinking water. Mice were additionally treated with weekly systemic DOS (0.05 mg/kg initial, followed by 1.5, 3, and 5 mg/kg). HIV burden among CD4^+^ T cells in blood was assessed by qPCR, 5 weeks after transfer.

To evaluate virological sanctuaries, 2 x 10^6^ CD45.1⁺ CD4⁺ T cells were HIV infected *in vitro* with empty backbone lentiviral vectors then transferred into CD45.2⁺ recipients. Recipient mice were vaccinated (or not) with FLUAD (1:20 of human dose) 18 hours later and treated weekly with systemic DOS as above. After 1, 3 or 12 months, CD45.1⁺ CD4⁺ T cells were recovered from lymphoid (lymph nodes, spleen) and non-lymphoid tissues (gut, brain) for quantification of integrated HIV DNA by qPCR, with similar HIV clearance results.

### Mathematical modelling

Integrated HIV DNA was quantified in CD4⁺ T cells from HIV⁺ individuals treated with DOS (dose escalation) for one week (*n* = 61). Residual proviral genomes were treated as rare events in a large cell population. Let *N*_cells_ denote the number of CD4⁺ T cells effectively interrogated per sample (2 x 10^5^) and *P*_cell_ the per-cell probability of carrying integrated HIV DNA. The expected number of proviruses per sample is:

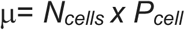

Under a Poisson sampling framework, the probability of detecting at least one provirus is

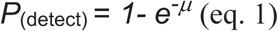

HIV-negative donor samples processed in parallel under identical conditions were used to define assay background and the effective single-event detection threshold. Accordingly, *P*_cell_ was conservatively _bounded_ by the experimentally defined assay detection limits (LOD 10^-12^; Ct≥ 40), using HIV negative controls in parallel. In the rare event regime relevant here, equivalent bounds are obtained under Poisson, binomial or hypergeometric sampling formulation; results are therefore independent of the specific model used.

To scale molecular frequency to the expected number of intact proviruses per individual, we defined:

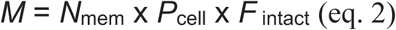

where *N*_mem_ denotes the resting memory CD4⁺ T cell pool and *F* _intact_ the fraction of proviral genetically intact proviruses.

Using conservative bounds anchored to assay detection limits as defined by HIV negative controls, this yields *M* < 1 expected intact provirus per individual under DOS treatment.

Time to viral rebound following ART withdrawal was modelled assuming a constant effective reactivation hazard *λ_eff_*, yielding a median rebound time:

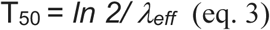

Upper-bound estimates assumed maximal reactivation competence of any residual intact proviral DNA. Functional competence was optionally parameterised as

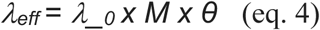

where *λ__0_* denotes the per-provirus annual reactivation rate and *θ* Ε (0,1] the fraction of intact proviruses capable of productive reactivation. Under these conservative assumptions, predicted rebound times extend beyond a human lifespan. *In vivo* analyses further supported long term viral clearance. By contrast, HIV+ samples not subjected to rejuvenation followed expected rebound kinetics (days) after antiretroviral interruption.

